# IFIT3 controls IFIT1 accumulation and specificity preventing self mRNA targeting during the innate immune response

**DOI:** 10.1101/2025.11.17.688928

**Authors:** Renata C Fleith, Xin Yun Leong, Taissa Ricciardi-Jorge, Harriet V Mears, Edward Emmott, Daniel S Mansur, Trevor R Sweeney

## Abstract

IFIT1 is among the highest expressed proteins following virus detection and interferon (IFN) induction and binds the 5’-end of ‘non-self’ viral cap0-mRNAs lacking 2’-O-methylation, blocking their translation. IFIT1 hetero-complexes with IFIT3, which enhances IFIT1 cap0-binding. Here, we demonstrate that IFIT3 is a master regulator of IFIT1, dictating its activity and stability. When expressed at high levels in the absence of IFIT3, IFIT1 can inhibit Semliki Forest virus replication, but can also inhibit translation of certain other ‘self’ IFN-stimulated genes (ISGs), including the important innate immune proteins ISG15 and IFITM1 as exemplars. However, IFIT1:IFIT3 complexing rescues ISG15 and IFITM1 from IFIT1 translation inhibition. We demonstrate that IFIT1 is degraded by the proteasome in the absence of IFIT3, but that direct binding to IFIT3 protects IFIT1 from degradation, ensuring IFIT1 accumulation only occurs along with IFIT3, demonstrating a mechanism to control the fine balance between antiviral activity and self-targeting in the face of unrelenting viral evolution.

## Introduction

The orchestration of the innate antiviral response relies on fine-tuning the expression and activation of sensor and effector proteins at the right time, place, and levels, to restrict virus infection without causing excessive tissue damage (1–3). Interferons (IFN) are cytokines which restrict viral infection and cell proliferation, by mediating the induction of hundreds of IFN-stimulated genes (ISGs). Although these genes have been intensively studied, our understanding of the relative importance of key effectors involved in controlling a given virus and how they interact, their dynamics of expression, and the regulation of their levels, in different cell types is still limited. These details have direct implications not only in understanding the antiviral response, but in many other physiological and pathological conditions where IFNs are implicated.

The family of IFN-stimulated proteins with tetratricopeptide repeats (IFITs) include some of the most highly expressed ISGs (4). Not only are they strongly upregulated following IFN stimulation, but their expression can be triggered directly by IFN regulatory factor 3 (IRF3) signalling immediately following pathogen sensing in cells (5, 6). Most mammals encode five IFIT paralogues: IFIT1, IFIT1B, IFIT2, IFIT3 and IFIT5 (7, 8). Due to differences in their promoter regions, their induction kinetics and expression levels can differ between IFIT genes and cell types (9–12). IFITs are both direct antiviral effectors and play roles in immune response modulation, apoptosis and inflammation (13–20). An increasing body of evidence associates IFITs to certain types of cancer and autoinflammatory diseases (21–23), both through poorly defined mechanisms.

IFIT1 functions by binding the 5’ cap of viral mRNAs lacking 2’-O-methylation (cap0-RNAs) with low nanomolar affinity and inhibits cap-dependent translation by blocking recruitment of the translation initiation complex (24–26). Since cellular mRNAs are typically further methylated on their first and sometimes second cap proximal nucleotides by cellular 2’-O-methyltransferases (2-O-MTases) producing cap1- and cap2-RNAs, respectively (27), current models report IFIT1 as distinguishing self from non-self RNAs. Consistent with its central role as a key ISG combating viral infection, IFIT1 was recently reported to be under strong selective pressure (28).

Viruses have evolved a variety of mechanisms to mimic cellular mRNAs and avoid IFIT1 restriction. Viruses that replicate in the nucleus, such as herpesviruses and retroviruses, hijack cellular enzymes to generate cap1-mRNAs (29, 30). Alternatively, orthomyxoviruses (e.g. influenza A virus (IAV)) and bunyaviruses ‘snatch’ pre-formed caps, by co-transcriptionally cleaving capped host RNAs and using them to drive viral translation (31). Viruses that replicate in the cytoplasm do not have access to the host transcriptional machinery. Instead, some cytoplasmic viruses, like Alphaviruses, have a stable RNA stem-loop at their 5’ terminus which precludes entry into the IFIT1 RNA binding channel (32). Others, including orthoflaviviruses (e.g. dengue, Zika, and West Nile viruses), coronaviruses, and poxviruses (e.g. vaccinia virus (VACV)) encode their own 2-O-MTases (33–38). Due to their importance for viral translation and immune evasion, viral capping enzymes and 2-O-MTases are an important focus of study for developing next generation vaccine targets, and several 2-O-MTase-deficient flavivirus and coronavirus vaccine candidates are attenuated in a IFIT1-dependent manner (26, 35, 39–43). IFIT1 was also found to bind to cap1-mRNA at micromolar affinity *in vitro*, which may overcome virally encoded 2-O-MTases. However, cap1-RNA binding raises the potential for host mRNA translation inhibition. These observations suggest that IFIT1 activity must be tightly regulated in cells to balance its role as an antiviral effector with its potential to cause cellular damage. How this regulation occurs is still largely unclear.

We and others have shown that members of the IFIT family form complexes *in vitro* and in cells and that an interaction between IFIT1 and IFIT3 enhances the cap0-mRNA binding affinity and antiviral activity of IFIT1 (44, 45). Here, we investigated whether this complex has further roles in IFIT1 regulation. Using knock-out (KO) cell lines and reconstitution experiments, we show that when expressed at high levels in isolation, IFIT1 inhibits the accumulation of canonically capped IFN-stimulated gene 15 (ISG15), a key innate immune regulator, and IFN-induced transmembrane protein 1 (IFITM1), an important Influenza A virus and SARS-CoV-2 restriction factor, following IFN treatment. Remarkably, this inhibition is reversed by IFIT1 complexing with IFIT3. In contrast, Semliki Forest Virus (SFV) replication was inhibited by high expression of IFIT1, irrespective of IFIT3 expression. Endogenous IFIT1 accumulation is dependent on IFIT3 binding following IFN stimulation and in the absence of IFIT1:IFIT3 hetero-oligomerisation, IFIT1 is degraded by the proteasome. Our findings reveal a new level of regulation within the host antiviral response allowing fine-tuning of specific antiviral proteins and enabling their elimination in situations where their expression may otherwise be detrimental to the cell.

## Results

### IFIT3 controls levels of endogenous IFN-induced IFIT1 in cells

We and others have previously observed that IFIT3 increases IFIT1 protein levels when exogenously co-expressed in cells (44, 45) by an unknown mechanism. We first investigated if endogenous IFIT3 expression also regulates endogenous IFIT1 in the IFN-induced antiviral cellular state. IFIT3 was depleted by siRNA knockdown (KD) in A549 or 293T cells before treatment with IFNα. Both cell lines have low basal IFIT1 expression, while A549s respond more robustly to IFN treatment.

IFIT3 KD greatly reduced both IFIT3 and IFIT1 protein levels in both A549 (Figure 1A) and 293T cells (Figure 1B). In contrast, IFIT2 KD did not affect IFIT1 protein expression (Supplementary Figure S1). To confirm this, we generated IFIT3 knockout (KO) 293T and A549 cells using CRISPR/Cas9 gene editing. Consistent with the KD experiments, both A549 and 293T IFIT3 KO cells showed lower IFIT1 levels upon IFN treatment when compared to wild-type (WT) cells (Figure 1C). In contrast, the levels of an unrelated ISG, PKR, were not changed by IFIT3 depletion (Figure 1C). IFIT1 mRNA level was unchanged in IFIT3 KO cells compared to WT cells (Figure 1D), confirming that IFIT1 expression was not affected at the transcriptional level following IFIT3 depletion.

**Figure 1.**
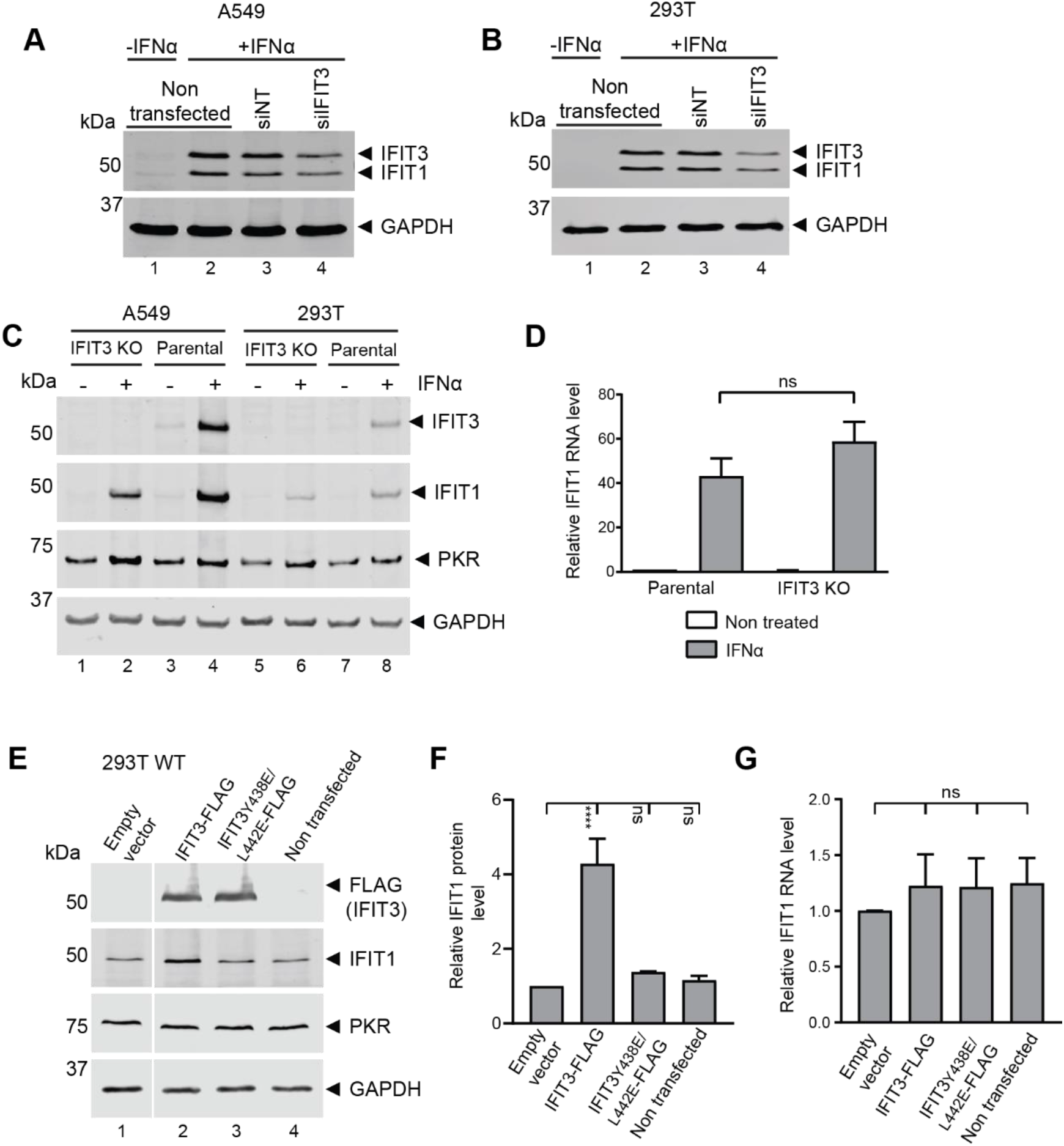
IFIT3 dictates IFIT1 protein level. A549 (A) and 293T (B) cells were transfected with pooled non-targeting siRNA (siNT) or siRNA targeting IFIT3 (siIFIT3). After 24 hours cells were treated with 100 U/mL of IFNα for a further 24 hours. Cell lysates were collected and analysed for immunoblotting 48 hours post transfection, probing with antibodies for the indicated proteins. C) A549 and 293T, IFIT3 KO and parental, cells were stimulated with 100 U/mL IFNα for 16 hours, then lysed and analysed by immunoblotting, probing with antibodies for the indicated proteins. A-C) GAPDH was included as loading control. D) Relative quantification of IFIT1 mRNA levels, in A549 IFIT3 KO cells stimulated with 100 U/mL IFNα for 16 hours, analysed by RT-qPCR, normalised by GAPDH expression and relative to non-stimulated parental condition. Data are mean +/-standard deviation (SD) of three independent experiments. Statistical analysis using unpaired, two-tailed students t-test comparing IFNα stimulated samples of parental and IFIT3 KO cells. ns=not significant. E, F, G) 293T cells were transfected with 0.33 pmol of empty vector or IFIT3-FLAG (wild type or Y438/L442E mutant) encoding plasmids. 24 hours after transfection, cells were treated with 100 U/mL IFNα for 16 hours. (E) Half the cells were lysed and analysed by immunoblotting probing with antibodies for the indicated proteins. (F) The graph shows the IFIT1 protein level normalized to GAPDH loading control, and relative to the empty vector condition. (G) The other half of the cells was used for RNA extraction and analysed by RT-qPCR. The graph shows IFIT1 RNA levels quantification, normalised by GAPDH expression and relative to the empty vector condition. (F and G) Data are mean +/-standard deviation (SD) of three independent experiments. Statistical analysis using one-way ANOVA. Asterisks indicate where IFIT1 level mean was significantly different (p < 0.05) between the empty vector and the other conditions as indicated. ns=not significant.

We next examined the effect of increasing IFIT3 expression on endogenous IFIT1 protein levels. WT 293T cells were transfected with WT IFIT3, or a Y438E/L442E mutant that we previously showed does not bind to IFIT1 (44), before cells were treated with IFNα. Exogenous expression of IFIT3 WT increased endogenous IFIT1 protein expression following IFNα treatment (Figure 1E-F), while IFIT1 mRNA expression was unaffected (Figure 1G). In contrast, overexpression of the IFIT3 Y438E/L442E mutant prior to IFNα treatment did not affect endogenous IFIT1 protein levels. Taken together these results show that endogenous IFIT1 protein expression is stabilised in the presence of IFIT3, this is dependent on their C-terminal interaction site and specific to the IFIT1:IFIT3 complex, since IFIT2 depletion did not affect IFIT1 expression levels. Moreover, cells have the capacity to produce more IFIT1 protein following IFNα treatment, but this level is strictly controlled in an IFIT3-dependent manner.

### IFIT1 is degraded by the proteasome in the absence of IFIT3

Given IFIT3’s essential role in controlling IFIT1 protein level (Figure 1) we next investigated how IFIT1 is degraded in the absence of IFIT3. IFIT3 KO cells were stimulated with IFNα for 16 hours, before incubation with fresh medium containing inhibitors for either the proteasomal or autophagy/lysosomal protein degradation pathways (Figure 2A).

**Figure 2.**
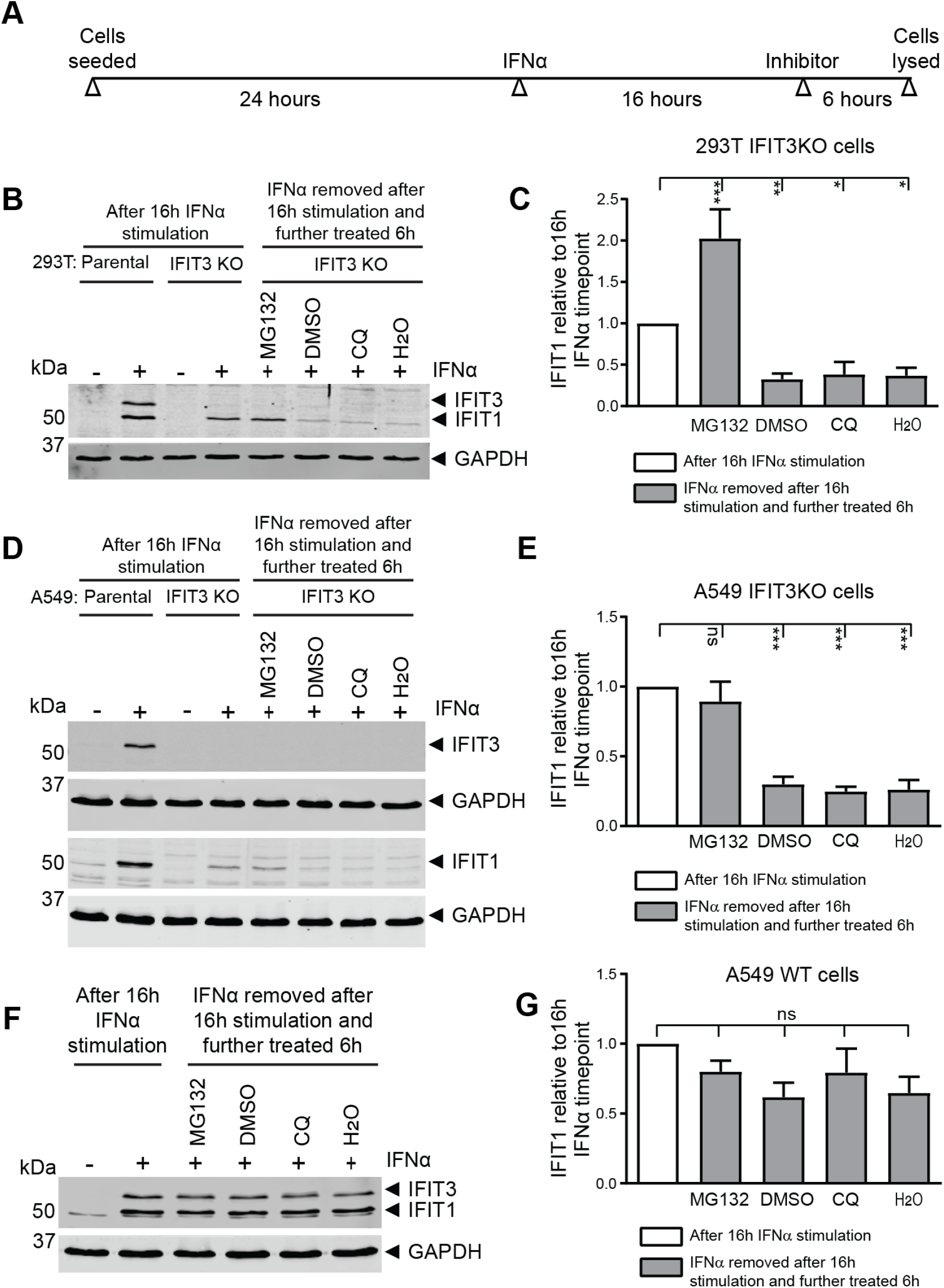
Endogenous IFIT1 is degraded in a proteasome-dependent manner in the absence of IFIT3. (A) Schematic representation of experiments performed in B-G. IFIT3 KO 293T (B/C), IFIT3 KO A549 cells (D/E), or wild type A549 (E/F) were stimulated with 100 U/mL IFNα for 16 hours, then washed twice, and incubated with fresh medium supplemented with indicated inhibitors or vehicle controls for further 6 hours, after which cell lysates were harvested for immunoblotting. In B and D wild type parental cell lysates were included as controls. GAPDH was included as loading control and membranes probed with antibodies for the indicated proteins. Quantification graphs (C, E and G) show IFIT1 protein levels normalised by GAPDH and relative to IFIT1 level in IFIT3 KO cell lysate (C and E) or IFIT1 level in wild type cell lysate (G) collected after 16hrs IFNα stimulation (white bars). Data is shown as mean +/-SD of three independent experiments. Statistical analysis using one-way ANOVA multiple comparison test. Asterisks indicate where IFIT1 level mean in inhibitors treated samples (grey bars) was significantly different (p < 0.05) to the levels before treatment (white bar). ns=not significant. CQ = chloroquine, DMSO = dimethyl sulfoxide. p <0.05, ** p <0.01, *** p <0.001.

In both 293T and A549 cells treated with water or DMSO (vehicle controls), IFIT1 levels decreased after IFNα removal, indicative of protein turnover (Figure 2B-E). Likewise, incubation with the autophagy-lysosomal inhibitor chloroquine (CQ) resulted in similarly decreased IFIT1 protein expression. However, treatment with the proteasomal inhibitor MG132 rescued IFIT1 levels at least to those at the point of IFNα removal, suggesting that endogenous IFIT1 is degraded by the proteasome. In MG132-treated 293T cells, we observed that IFIT1 levels were ∼2-fold higher following inhibitor treatment than they were immediately following IFNα removal, indicative of further protein synthesis from IFIT1 mRNA (Figure 2B-C). Similar results were observed with another proteasome inhibitor (Lactacystin), which rescued IFIT1 protein expression, and further autophagy-lysosome inhibitors (bafilomycin A1 and 3-methyladenine) which failed to rescue IFIT1 (Supplementary Figure S2).

Finally, we observed increased expression of ectopic IFIT1 in 293T cells treated with MG132, but not with autophagy-lysosome inhibitors (Supplementary Figure S3). The levels of IFIT2, IFIT3 or IFIT5 proteins was comparable between treated and untreated cells (Supplementary Figure S3). Therefore, in the absence of IFIT3, in both endogenous and overexpression systems, IFIT1 is turned over in a proteasome-dependent manner. In contrast, in WT A549 cells, where IFIT3 is also expressed, IFIT1 levels were comparable between inhibitor and vehicle treatments, demonstrating that in the presence of IFIT3, IFIT1 is protected from proteasomal degradation (Figure 2F and G).

### Direct IFIT3 binding prevents IFIT1 degradation

We next investigated if direct interaction between IFIT1 and IFIT3 was required for protection of IFIT1 from proteasomal degradation. We trans-complemented 293T IFIT3 KO cells with IFIT3, prior to IFNα and MG132 treatment (Figure 3A, and 3B top panel). We observed increased endogenous IFIT1 expression in cells ectopically expressing WT IFIT3 compared to the empty vector control, but not in cells expressing the IFIT3 Y438E/L442E mutant. Likewise, MG132 treatment rescued IFIT1 expression in empty vector or IFIT3 Y438E/L442E mutant-expressing cells, but not in WT IFIT3-expressing cells (Figure 3B-C).

**Figure 3.**
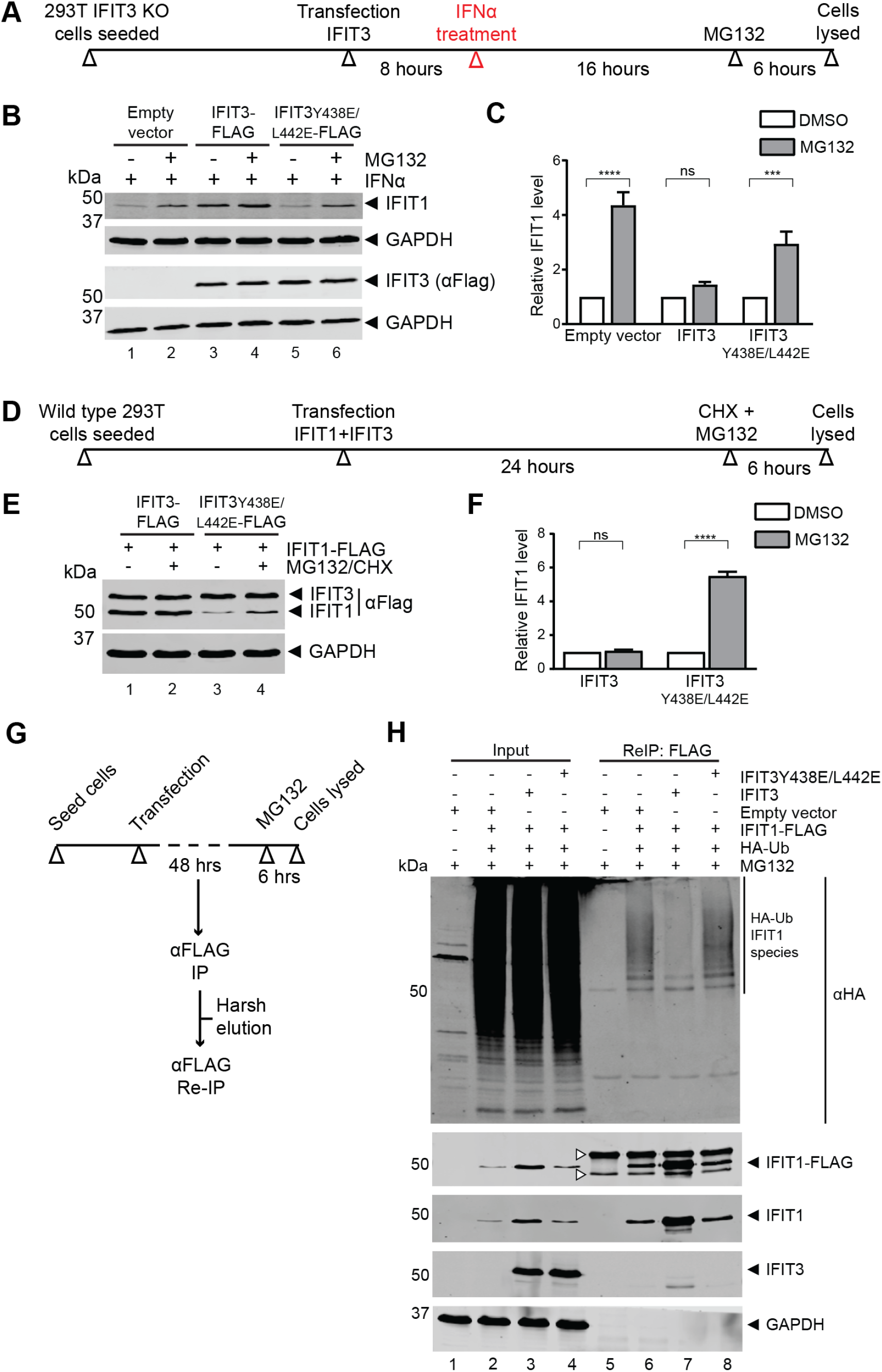
IFIT3 interaction prevents IFIT1 proteasomal degradation. (A-C) 293T IFIT3 KO cells were transfected with equimolar amounts (0.33 pmol) of empty vector or FLAG-tagged wild type or Y438E/L442E mutant IFIT3 encoding plasmids for 8 hours, then stimulated with 100 U/mL IFNα for 16 hours. Subsequently, cells were washed twice and further incubated with media containing DMSO or MG132 for 6 hours. Cell lysates were analysed by immunoblot using anti IFIT1 or anti FLAG, and anti GAPDH antibodies. (D-F) Wild type 293T cells were co-transfected with 1.5 µg FLAG-tagged pcDNA IFIT1 plus 2.5 µg FLAG-tagged wt or Y438E/L442E mutant pcDNA IFIT3 encoding plasmids for 24 hours and further incubated in medium containing cycloheximide (CHX) plus MG132 or DMSO for 6 hours. Cell lysates were harvested for analysis by immunoblot using antibodies anti FLAG and GAPDH (probed as loading control). GAPDH was used as loading control in both blots. (C, F) Quantification graph shows (C) endogenous IFIT1 or (F) IFIT1-FLAG protein levels normalized to GAPDH and relative to the DMSO control of each transfected condition. Data shown as mean +/-SD of three independent experiments. Statistical analysis using (C) one-way ANOVA or (F) unpaired, two-tailed students t-test, comparing the DMSO and MG132 of each transfection condition. Asterisks indicate where IFIT1 level mean in MG132 treated samples (grey bars) was significantly different (p < 0.05) to the DMSO controls (white bars). ns=not significant. (G, H) Overexpressed IFIT1 is directly HA-ubiquitinated in the absence of IFIT3. As outlined in G, 293T cells were transfected with the indicated protein encoding plasmids. 48 hours post-transfection, cells were incubated with fresh medium containing MG132 for 6 hours, after which cell lysates were harvested. IFIT1-FLAG was pulled down in two rounds of immunoprecipitation (IP and ReIP). Initial lysate inputs and ReIP eluates were analysed by immunoblotting using the antibodies for the indicated proteins. Arrows indicate non-specific bands. GAPDH was probed as loading control for input and equal volume of eluates was loaded for IP lanes.

We then co-transfected ectopic IFIT1 into 293T cells with WT or Y438E/L442E mutant IFIT3, before treatment with cycloheximide, to prevent new protein synthesis, and MG132 (Figure 3D, and E top panel). When WT IFIT3 was co-expressed with IFIT1, MG132 treatment had no effect on IFIT1 protein levels. However, in the presence of the Y438E/L442E mutant of IFIT3, MG132 rescued IFIT1 expression, albeit to lower levels compared to co-expression with WT IFIT3 (Figure 3E-F). Taken together, these data show that direct interaction between IFIT1 and IFIT3 is essential to protect IFIT1 from degradation by the proteasome.

As we observed that the 293T overexpression system recapitulated endogenous IFIT1 proteasome-dependent degradation, we co-transfected these cells with plasmids encoding HA-tagged ubiquitin, FLAG-tagged IFIT1, non-tagged IFIT3 or empty vector, before treatment with MG132. IFIT1-FLAG was purified from cell lysates in two rounds of anti-FLAG immunoprecipitation (Re-IP), and the samples analysed by immunoblot (Figure 3G). A stringent wash buffer was used to remove proteins co-precipitating with IFIT1, as indicated by the depletion of the IFIT3 signal, to ensure the anti-HA signal could be reliably attributed to IFIT1 (Figure 3H). Consistently, we observed minimal anti-HA signal in eluates lacking IFIT1-FLAG following Re-IP (Figure 3H, lane 5).

HA-ubiquitin signal was observed in Re-IP eluates, ranging between 50-250 kDa when IFIT1-FLAG (56 kDa) was expressed in isolation (Figure 3H, lane 6). When IFIT3 WT was co-expressed with IFIT1, HA-ubiquitin signal was greatly reduced, despite higher overall IFIT1 expression (Figure 3H, compare lanes 6 and 7). In contrast, co-expression of the IFIT3 Y438E/L442E mutant failed to protect IFIT1 from ubiquitination (Figure 3H, compare lanes 7 and 8). These results show that IFIT1 is ubiquitinated through a mechanism regulated by direct interaction with IFIT3.

### IFIT1 is sufficient to restrict SFV when expressed at high levels

In an ISG‐targeted CRISPR‐Cas9 KO screen, IFIT1 and IFIT3 were identified as dominant antiviral effectors against certain alphaviruses (46), while IFIT3 co-expression was further reported to enhance alphavirus restriction by IFIT1 (45). Alphaviruses possess a single-stranded positive-sense RNA genome that encodes a capping enzyme but lacks 2-O-MTase activity, resulting in a ‘non-self’ 5’ cap0 structure. Having demonstrated that direct binding of IFIT3 controls endogenous IFIT1 protein level through protection from proteasomal degradation, we examined the effect of high IFIT1 expression on alphavirus infection in the absence of IFIT3.

To address this, we further knocked out IFIT1 from our IFIT3 KO cells (Figure 4A) to generate IFIT1/3 double KO cells. To force IFIT1 accumulation besides faster decay in the absence of IFIT3, we designed a new system where IFIT1 WT or with a mutation at R38 that is essential for cap0-mRNA binding (24) was cloned into a plasmid under the control of the strong eukaryotic translation elongation factor 1A (eEF1A) promoter (Figure 4B). For comparison, we used the same system to co-express both IFIT1 and IFIT3 (WT or Y438/L442E mutant cannot bind IFIT1) separated by a 2A ribosomal stop-go peptide sequence from a single plasmid, ensuring efficient co-expression of both proteins in transfected cells (Figure 4B). Unlike previous studies, this strategy enabled assessment of IFIT1 activity when expressed at high levels in the absence of IFIT3 (Figure 4C).

**Figure 4.**
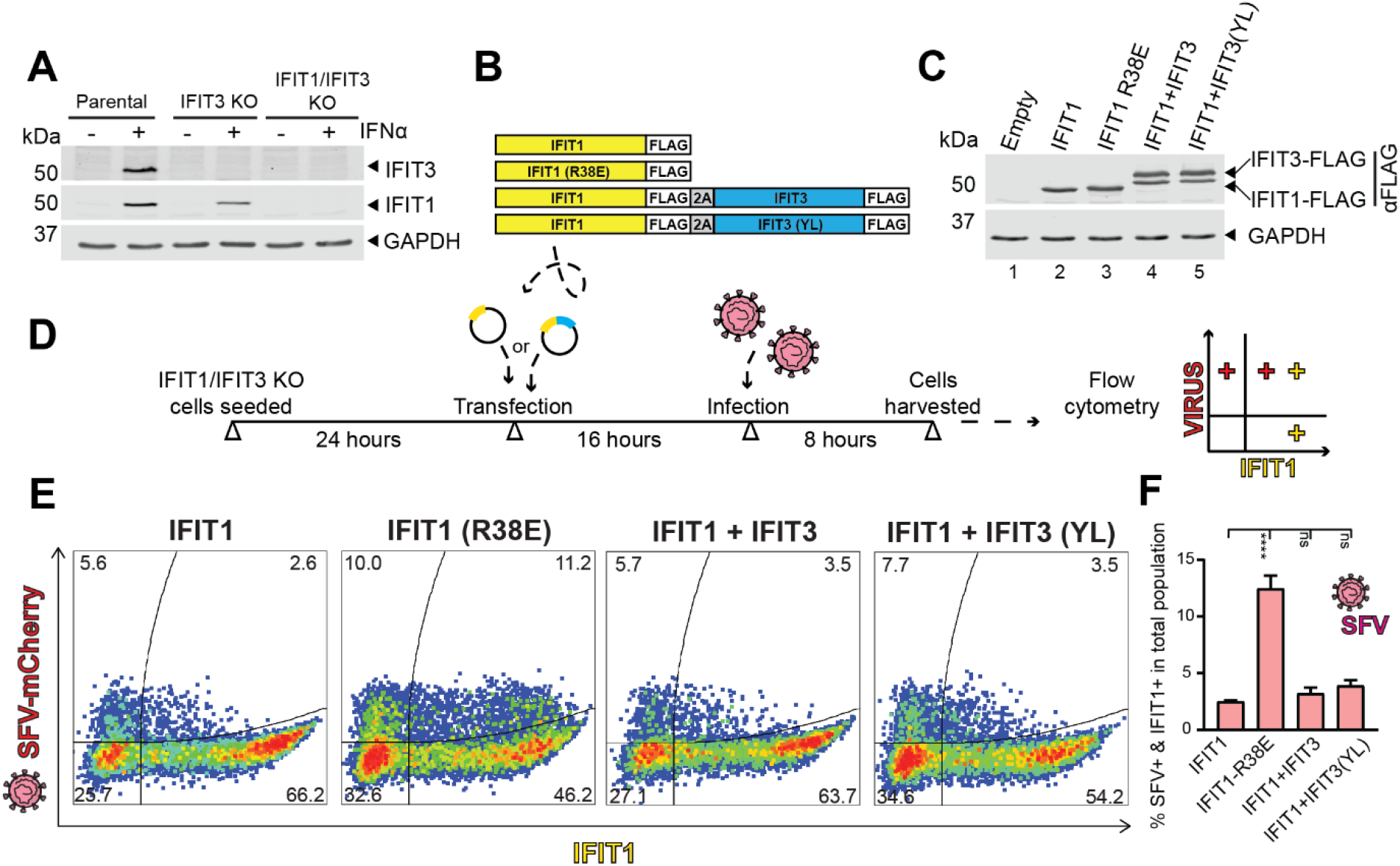
IFIT1 restricts SFV infection. A) Parental, IFIT3 KO and IFIT1/IFIT3 double KO 293T cells were stimulated with IFNα for 24 hours. Cell lysates were harvested for analysis by immunoblot using antibodies anti IFIT1, IFIT3 and GAPDH (probed as loading control). B) Schematic of IFIT1-FLAG and IFIT3-FLAG gene inserts into pTwist EF1 Alpha. In dual expression constructs IFIT genes were separated by a 2A ribosomal stop-go peptide sequence. C) IFIT1/IFIT3 double KO 293T cells were transfected with 0.4 pmol of each construct showed in B, then harvested after 16 hours to compare IFIT1 expression levels. Lysates were analysed by immunoblot using antibodies anti FLAG and GAPDH (probed as loading control). D) Schematic of experiment shown in E. Cells were transfected with 0.4 pmol of each plasmid and after 16 hours infected with SFV-mCherry at MOI 2.5. After 8 hours infection, cells were harvested, fixed, and stained with antibody anti IFIT1. The percentage of double IFIT1 and virus positive cells was determined by flow cytometry, with gates defined based on empty vector transfected, non-treated cells (Figure S4). Data are representative flow cytometry dot plots of three experiments with percentage of cells in each quadrant shown. F) Quantification of the percentage of double IFIT1 and virus positive cells in the total population. Data are mean +/-standard deviation (SD) of three independent experiments. Statistical analysis using one-way ANOVA. Asterisks indicate where the percentage of double positive cells was significantly different (p < 0.05) between the IFIT1 WT and the other conditions as indicated. ns=not significant.

Therefore, we used the IFIT1/IFIT3 double KO 293T cells and overexpression system to first examine the effect of IFIT1 alone or in the presence of IFIT3 on replication of Semliki Forest virus (SFV), chosen as a model alphavirus with greater inherent resistance to IFIT1 ((47). We infected reconstituted IFIT1/3 double KO cells with an SFV mCherry reporter virus and analysed cells after 8 hours for IFIT1 and SFV-mCherry expression using flow cytometry (Figure 4D-F). In cells expressing the RNA-binding mutant IFIT1 R38E ∼12.5% of cells were positive for both IFIT1 and mCherry-SFV (Figure 4F). In contrast, expression of wildtype IFIT1 reduced the number of double-positive cells to ∼2.5%, in-line with previous data that IFIT1 inhibits SFV infection in an RNA-binding-dependent manner (46, 47). Surprisingly, IFIT1-mediated SFV restriction was comparable when IFIT1 was expressed alone or co-expressed with WT or the YL mutant of IFIT3 demonstrating that when expressed to high levels, IFIT1 in isolation can inhibit SFV replication, overcoming the need for IFIT3 to increase its cap0 affinity.

### IFIT3 protects self mRNA from IFIT1 translation inhibition

We next asked why this complex regulatory relationship between IFIT1 and IFIT3 has evolved. Given IFIT1’s potency as a translation repressor (20) and IFIT3’s reported capacity to enhance IFIT1 cap0, but not cap1 viral mRNA binding, we hypothesised that IFIT3 may affect IFIT1 interaction with ‘self’ mRNAs, preventing unintended host translation inhibition. As IFIT1 is typically expressed at very low levels in uninfected cells and IFIT1’s predominant role as an ISG we examined its effect in IFN-stimulated cells. Both ISG15 and IFITM1 were previously reported as sensitive to IFIT1 translation inhibition in cap methyl transferase 1 depleted cells where mRNAs lacked 2’-O-methylation (13). Like IFIT1, ISG15 and IFITM1 are strongly induced at the mRNA and protein level in response to IFN stimulation. We therefore examined the effect of high levels of IFIT1 on ISG15 and IFITM1 protein levels. For that, we first overexpressed IFIT1 WT or R38E mutant in IFIT1/3 double KO cells under the control of the EF1a promoter to achieve high levels of the protein in the absence of IFIT3. Cells were then treated with interferon for 24 hours and the lysates analysed by immunoblotting. While ISG15 and IFITM1 levels were not impacted by IFIT1-R38E RNA binding mutant, the overexpression of the WT protein remarkedly reduced their levels (Figure 5A). These data show for the first time that translation of endogenously expressed and canonically capped ISG mRNA is inhibited by IFIT1, and that this inhibition is through IFIT1’s RNA cap-binding activity. In the other hand, the expression of PKR, another ISG, was not impacted for IFIT1, suggesting that this effect is specific for some self-RNAs (Figure 5A).

**Figure 5.**
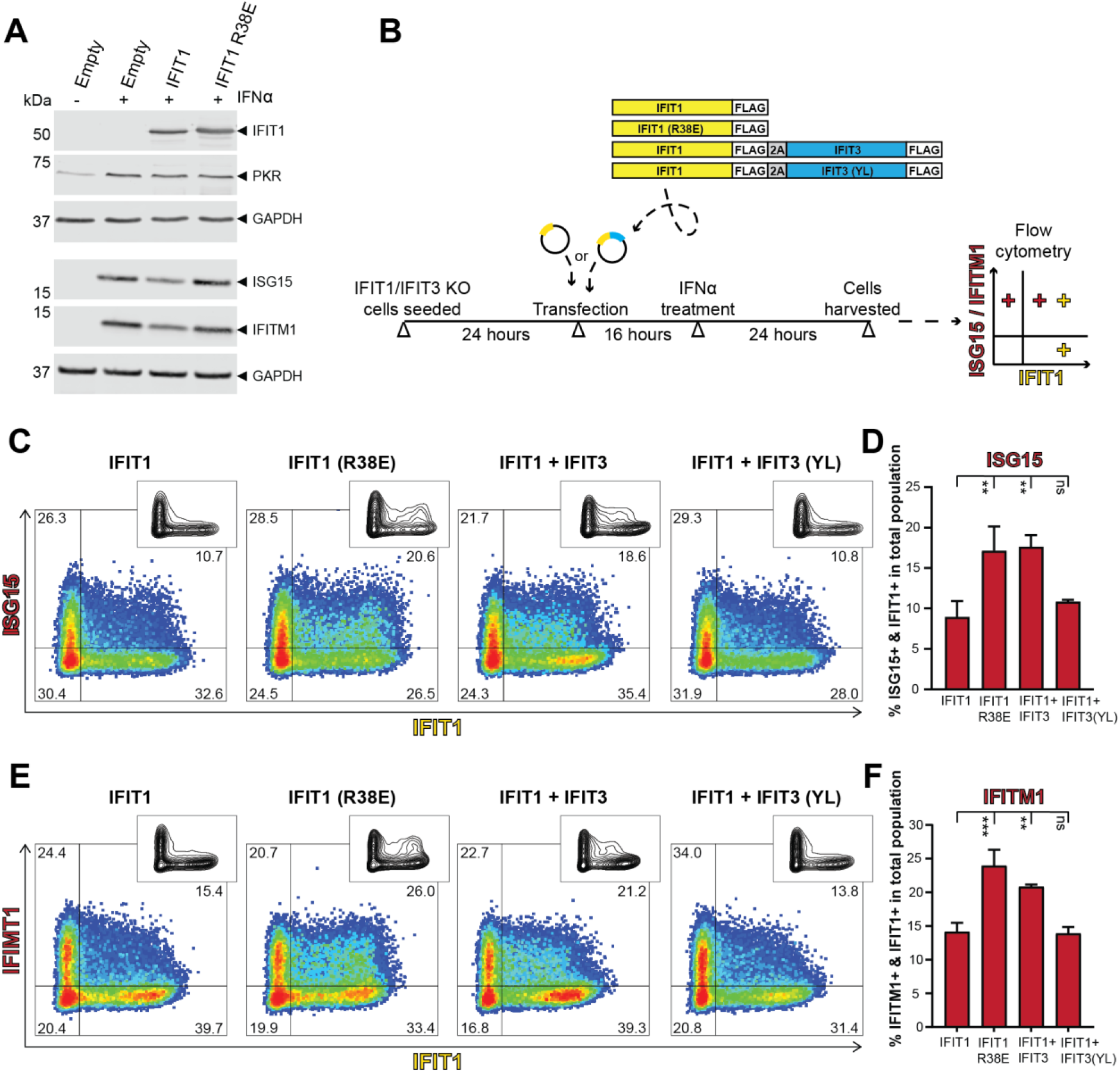
IFIT3 protects ISG15 and IFITM1 mRNA from IFIT1 translation inhibition. A) 293T IFIT1/IFIT3 double KO cells were transfected with 0.4 pmol of plasmids expressing IFIT1 or IFIT1-R38E mutant under the control of an EF1α promoter. 16 hours after transfection, cells were treated with 500 U of IFNα. Lysates were analysed by immunoblotting 24 hours after treatment. GAPDH was included as loading control and membranes probed with antibodies for the indicated proteins. B) Schematic of experiment shown in C-F. Cells were transfected with 0.4 pmol of each IFIT plasmid under the control of an EF1α promoter and after 16 hours treated with 500 U of IFNα (F,G). After 24 hours IFNα treatment, cells were harvested, fixed, and stained with antibodies. The percentage of double IFIT1 and ISG15 positive (C), or IFIT1 and IFITM1 positive (E) cells was determined by flow cytometry, with gates defined based on empty vector-transfected, non-treated cells. Data are representative flow cytometry dot plots of three experiments, with the percentage of cells in each quadrant shown. Corresponding contour plots are also shown in C and E for ease of comparison. Quantification of the percentage of double IFIT1 and ISG15 or IFITM1 positive cells in the total population is shown in D or F respectively. Data are mean +/-standard deviation (SD) of three independent experiments. Statistical analysis using one-way ANOVA. Asterisks indicate where the percentage of double positive cells was significantly different (p < 0.05) between the IFIT1 WT and the other conditions as indicated. ns=not significant.

Then we assessed the impact of IFIT3 co-expression in IFIT1-self RNA translation inhibition at cell level. IFIT1/3 double KO cells were transfected with IFIT1 or IFIT1/IFIT3-expressing plasmids to enable high expression of both proteins before treatment with IFN and analysis by flow cytometry (scheme shown in Figure 5B). Similar percentages of IFIT1-positive cells were detected after transfection of IFIT1, IFIT1 R38E and IFIT1/IFIT3 Y438/L442E plasmids (Figure S5). As shown in Figure 5C-F, a significantly higher proportion of IFIT1 expressing cells were positive for ISG15 and IFITM1, respectively, when cells were transfected with the RNA-binding mutant, IFIT1 R38E, compared to WT IFIT1.

In contrast, both ISG15 and IFITM1 protein levels were restored when IFIT3 was co-expressed with IFIT1. This protection was dependent on direct interaction between IFIT1 and IFIT3 as the IFIT3 Y438/L442E mutant failed to restore ISG15 or IFITM1 protein levels (Figure 5C-F). Therefore, direct binding of IFIT3 to IFIT1 protected ISG15 and IFITM1 from IFIT1-dependent translation inhibition. Together, our data reveal that IFIT1 can repress the translation of host ISG mRNAs that could reduce the efficacy of the innate immune response to respond to infection but that this is tempered by IFIT3. Therefore, IFIT3 both enables IFIT1 accumulation to the expression level required for anti-viral activity but prevents IFIT1 inhibition of self mRNAs.

## Discussion

During the antiviral response, hundreds of ISGs are upregulated. Their expression is not uniform, and cells have evolved mechanisms to precisely control the duration and levels of expression of individual ISGs (48, 49). IFIT1 is among the earliest expressed ISGs and is detectable even before type-I IFN (50). However, cells do not normally express IFIT1 at basal levels (6, 50, 51), suggesting tight regulation under resting conditions. Moreover, despite its high expression in response to IFN, we and others have shown that human IFIT1 expresses poorly in previously used ectopic overexpression systems compared to other IFIT family members, suggesting that cells have developed mechanisms to specifically regulate IFIT1 protein levels (44, 52). We have previously shown that IFIT3 is a key co-factor to enhance IFIT1 RNA binding and protein stability both *in vitro* and in cells. In this study, using a plasmid promoter system that drives strong expression of IFIT1 so that it accumulates to high levels in the absence of IFIT3, we demonstrate that IFIT1 alone inhibits replication of a cap0 RNA virus, SFV, but also impacts translation of host proteins.

We and others have previously shown that IFIT3 increases IFIT1 affinity to cap0 RNA, being particularly important for binding of viral RNAs with structured 5’UTRs, such as modified orthoflaviviruses vaccine candidates lacking 2’-O methylation (44, 45). However, co-expression of IFIT3 was also reported to enhance IFIT1-dependent inhibition of Venezuelan equine encephalitis virus (VEEV) strain TC83, an alphavirus lacking stable structure at the 5’ end (45). IFIT1 and IFIT3 were also reported as key restriction factors for VEEV in a human cell model system (46), while, in work performed in a murine model system, different alphaviruses were reported to have varying sensitivity to Ifit1 restriction due to differing stability of 5’ RNA secondary structure (47). However, here we show that in human cells, high levels of IFIT1 restricts alphavirus SFV replication even in the absence of IFIT3, and addition of IFIT3 does not further inhibit replication. Our data therefore support a revised model for IFIT1:IFIT3 complexing in antiviral activity; the main role of IFIT3 is maintaining IFIT1 at sufficiently high levels to perform its antiviral activity, consistent with IFIT3 being a key restriction factor along with IFIT1 (46), while concomitantly protecting susceptible host mRNAs from IFIT1 translation inhibition (Figure 6). McDougal et al. (28) recently showed that overexpression of IFIT1 from different species had varying effects on replication of the alphavirus VEEV. Interestingly, the different IFIT1 proteins had very different levels of overexpression in human Huh7.5 cells. It is possible that the proteins from different species have varying susceptibility to degradation in human cells, however, co-expression of complementary species-specific IFIT3 or potential for interaction with human IFIT3 was not assessed.

**Figure 6.**
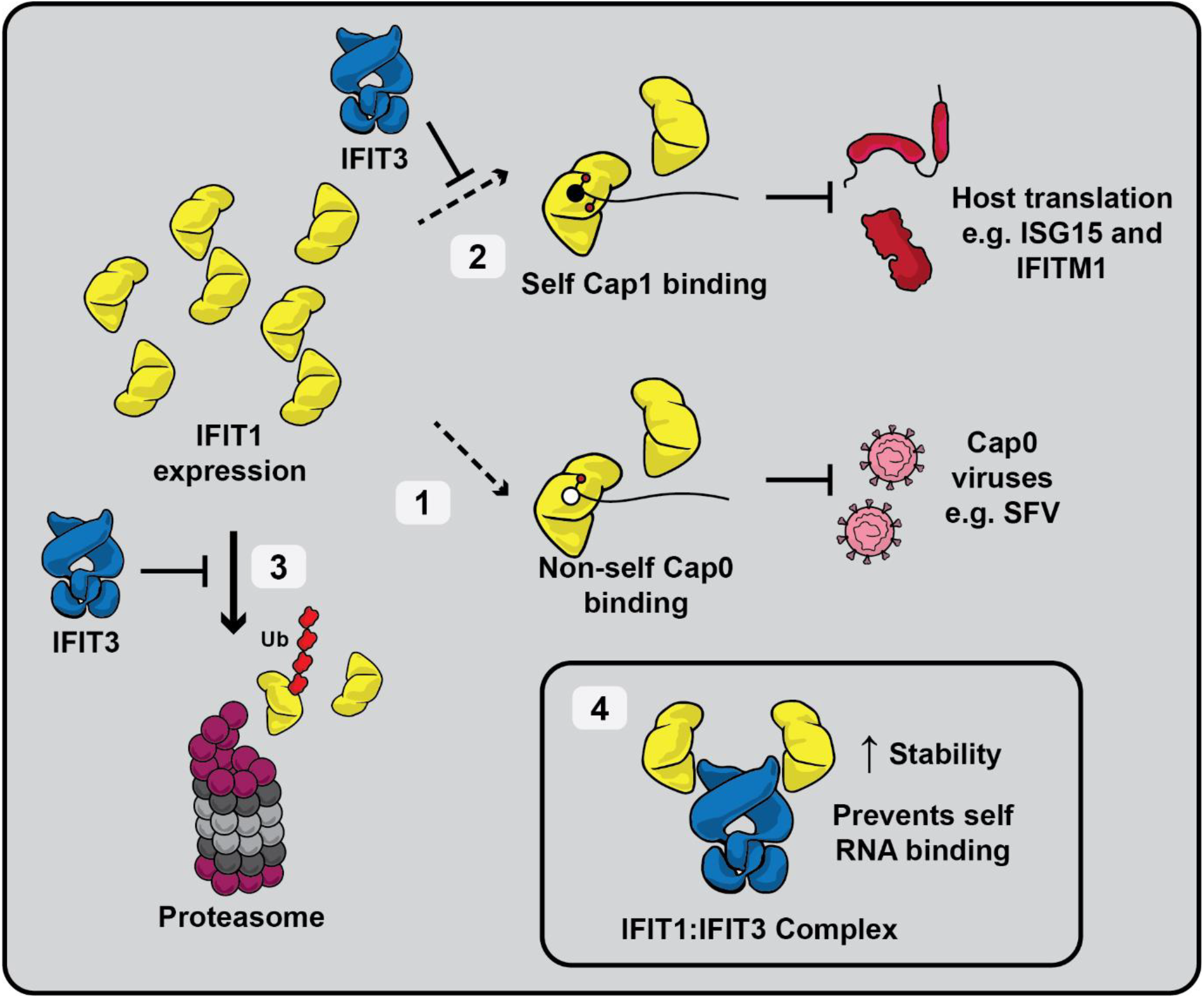
Model for the evolutionary role of IFIT1:IFIT3 complexing. **1)** When expressed at high levels in isolation, IFIT1 (yellow) can restrict viruses with cap0 mRNAs through its RNA-binding activity, as shown here for SFV. **2)** However, in these conditions, IFIT1 also inhibits susceptible canonically-capped self-RNAs, including of innate immune molecules, such as ISG15 and IFITM1. Co-expression of IFIT3 (blue) prevents IFIT1-mediated host-translation inhibition. **3)** When expressed in isolation, IFIT1 can be ubiquitinated in cells and rapidly degraded by the proteasome, which is prevented by co-expression of IFIT3. **4)** IFIT1:IFIT3 complex formation prevents both IFIT1 degradation, and IFIT1 binding to self-RNAs. This mechanism allows cells to accumulate high levels of IFIT1, ensuring robust antiviral activity towards cap0 RNA viruses, without a detrimental impact in host translation. Cap0 and cap1 RNA are depicted by an open and a closed circle, respectively. Ub = ubiquitin.

There may be different reasons why cells need to control IFIT1 levels. For example, here, we show that IFIT1 can inhibit the translation of ISG15, a key innate immune regulatory factor. ISG15 is a small ubiquitin-like protein that is covalently linked to certain host and viral proteins that impacts replication of a range of viral families through a variety of strategies (reviewed in (53, 54). Disruption of ISG15 translation by aberrant IFIT1 expression would therefore diminish the cell’s ability to respond to a wide variety of viral challenges. IFIT1 also inhibited translation of IFITM1, a membrane-associated ISG with broadly documented antiviral activity against viruses including HIV, SARS-CoV-2 and influenza (55). IFIT1 has been further reported to downregulate translation of other host ISGs when the host cap1 methyltransferase, CMTR1, was depleted (13, 56), increasing the number of host mRNAs that may be inhibited by uncontrolled IFIT1 expression. Interestingly, IFIT3 was identified as an ISG that was resistant to IFIT1 translation inhibition after CMTR1 depletion (13). IFIT3 resistance to IFIT1 translation inhibition could provide a failsafe to ensure expression of IFIT3 to modulate IFIT1 stability and activity. Importantly, CMTR1 is itself an ISG, suggesting that upregulation of CMTR1 is necessary to fortify cellular mRNAs in the presence of high IFIT1 expression.

Independent of its role in translation regulation, IFIT1 is involved in multiple cellular signalling pathways (13–20). Dysregulated IFIT1 expression has therefore been associated with certain interferonopathies, autoimmune and inflammatory conditions. IFIT1 is also in the group of ISGs with IFN-related damage resistance signature (IRDS), which are linked to malignancy progression and radio and chemotherapy resistance in multiple cancers (21, 57). In oral squamous cancer for instance, overexpression of IFIT1 and IFIT3 is associated with tumour growth and metastasis by enhancing recycling of phosphorylated epidermal growth factor receptor (58). Therefore, IFIT1 regulation may be necessary in cellular homeostasis. We found that IFIT1 is ubiquitinated and rapidly turned over in a proteasome-dependent manner. Strikingly, in the presence of IFIT3, IFIT1 ubiquitination and degradation were dramatically reduced. We previously showed that a highly conserved YxxxL motif in the C-terminal region of both IFIT1 and IFIT3 is essential for their interaction (44). Mutating the YxxxL motif in IFIT3 eliminated its ability to stabilise IFIT1 revealing that direct binding between the two proteins is essential to protect IFIT1 from ubiquitination and degradation.

Therefore, while controlling IFIT1 levels and specificity to cap0-RNA through the regulatory role of IFIT3 can mitigate potentially detrimental effects of self RNA binding, it may also reduce the antiviral scope of IFIT1. This is especially true for numerous viruses that have evolved mechanisms to generate a cap1 structure to mimic that of the host (19). This highlights the difficult balance in maintaining and evolving robust antiviral mechanisms in the face of rapidly evolving pathogens. Indeed, while IFITs are as old as the IFN system (7), a recent study shows that IFIT1 is under strong evolutionary pressure and rapidly evolving (28). Notably, in mice and some other rodents, the IFIT1 gene orthologue and the C-terminal region of *Ifit*3, including the YxxxL motif, have been lost (8). Instead, the closely related *Ifit1b* has been duplicated twice, yielding three paralogues: *Ifit1, Ifit1b*, and *Ifit1c*. Indeed, we have previously reported that while murine *Ifit1* has strong cap0-RNA-binding selectivity, *Ifit1b* preferentially binds to cap1-RNA and *Ifit1c* acts as a stimulatory cofactor for both *Ifit1* and *Ifit1b*, enhancing expression and translation inhibition activity analogous to the role of human IFIT3 (59).

Together, our findings reveal the evolution of an elegant mechanism to fine tune IFIT1 levels and activity by the immune system to reduce targeting of self mRNAs by expression of another, related ISG, protein, IFIT3. Our data expand our understanding of the complex interplay of members of the IFIT family, which are a fundamental component of mammalian innate immunity, balancing potent antiviral activities with an imperative to maintain cellular health.

## Materials and Methods

### Cells

Human embryonic kidney cells (293T) and adenocarcinomic alveolar basal epithelial cells (A549) (both from European Collection of Authenticated Cell Cultures) were maintained in high glucose (4500 mg/L) Dulbecco modified Eagle medium (DMEM) (Sigma-Aldrich) supplemented with 10% foetal bovine serum (FBS), penicillin (100 U/mL), streptomycin (100 μg/mL) and L-glutamine (2 mM) (all Thermo Fisher Scientific, TFS) and incubated at 37 °C with 5% CO_2_. Cells were seeded in medium without antibiotics before transfection. IFIT3 knockout cells were generated by CRISPR gene editing, adapting a plasmid-based protocol previously described (60). Briefly, annealed oligonucleotides for sgRNA ACACCTAGATGGTAACAACG (61) were cloned between BbsI restriction sites into the PX458 plasmid (gift from Feng Zhang, Addgene plasmid #48138) encoding for Cas9 and EGFP. The plasmid was transfected in cells using Lipofectamine 2000 (TFS). After transfection, single EGFP-positive cells were fluorescence-activated cell sorted for clonal cell generation. Empty vector transfected cells were sorted as parental cells. IFIT1/IFIT3 doble KO 293T cells were generated by knocking out the IFIT1 gene in the IFIT3 KO background using a lentivirus-based system as previously described (62). Briefly, for lentivirus production four annealed oligonucleotides for IFIT1 (caccgAGGCATTTCATCGTCATCAA, caccgTACAAATGGTGATGATCATC, caccgCTAGACACCAAATACAGTGT, caccgCAGCTTCTTTTAAGCTCTTC) were inserted into the lentiCRISPRv2-Blast plasmid (gift from Mohan Babu, Addgene plasmid # 83480) between BsmBI restriction sites, and along with pMD2.G and psPAX2 plasmids (gifts from Didier Trono, Addgene plasmids #12259 and #12260 respectively). Cells were transduced and antibiotic selected for one week and sorted for clonal cell generation. Both single and double KO cell lines were analysed by immunoblotting after IFN induction and validated by Sanger sequencing.

### IFN treatment

Cells were treated with medium containing 100 U/mL IFNα 2a (PBL Assay Science) for induction of endogenous IFITs, and 500U/mL for induction of ISG15 and IFITM1, then incubated for the time indicated in each figure.

### Plasmids

IFIT constructs possessing a CMV promoter in pcDNA3.1 were previously described (44). IFIT3 Y438E/L442E mutant and Non-FLAG IFIT3 were generated by PCR site-directed mutagenesis using the wild type IFIT3 construct as template. All constructs were verified by Sanger or whole plasmid sequencing (Genewiz). For figures 4 and 5, FLAG-tagged IFIT genes were custom synthesised and cloned in a pTwist vector containing Human Elongation Factor Alpha (EF1 Alpha) promoter (Twist Bioscience). Dual expression vectors were designed encoding both IFIT1-FLAG and IFIT3-FLAG separated by a Thosea asigna virus (T2A) ribosomal stop-go peptide sequence (EGRGSLLTCGDVEENPGP). Both IFIT1 and IFIT3 mutants in this vector were generated by PCR site-directed mutagenesis.

### Transfection

Cells were seeded in antibiotic-free medium and transfected the next day using Opti-MEM reduced serum medium (TFS) and Lipofectamine 2000 (TFS) according to the manufacturer instructions. For plasmid DNA transfection a ratio of 2 μL of Lipofectamine 2000 was used for each μg of DNA (see figure legends for amount of plasmid in each experiment, or below for ReIP). For siRNA transfection, 2 μL of Lipofectamine 2000 was used for 40 pmol of siRNA in a 12-well plate format. All siRNAs used were predesigned siGENOME in the SMARTpool format (Dharmacon/Horizon Discovery).

### RNA extraction and RT-qPCR

Detached cells were washed twice with 1X phosphate-buffered saline (PBS - 137 mM NaCl, 2.7 mM KCl, 10 mM Na2HPO4, and 1.8 mM KH2PO4) and the RNA purified with TRIzol (TFS). RNA samples were treated with DNaseI (TFS), and reverse transcribed with Moloney Murine Leukaemia Virus Reverse Transcriptase (Promega) using random hexamers (TFS). qPCR reactions were prepared with cDNA, primers for GAPDH (63) and IFIT1 (64) and PowerUp SYBR master mix (TFS) using the manufacturer’s recommended parameters. Relative quantification was expressed as 2-ΔΔCt, normalised to GAPDH expression, with non-stimulated parental conditions (Figure 1D) or empty vector (Figure 1G) used as calibrator.

### Proteasome and autophagy-lysosome inhibition assays

For assays using the endogenous system, cells were stimulated with IFNα 2a for 16 hours then rinsed twice with PBS, and further incubated with fresh medium supplemented with inhibitors of autophagy-lysosomal pathway or proteasome pathway or their vehicle controls for 6 hours: MG132 (Selleckchem) 10 µM, Lactacystin (Enzo Life Sciences) 20 µM, Chloroquine (Sigma-Aldrich) 100 µM, 3-methyladenine (TFS) 5 mM and Bafilomycin A1 (Selleckchem) 100 nM. For the overexpression system, different FLAG-tagged IFITs encoding pcDNA3.1 plasmids were transfected into wild type 293T cells for 24 hours then further incubated with fresh medium containing cycloheximide (100 µg/mL) (Sigma-Aldrich) plus proteasome or lysosome inhibitors for 6 hours.

### Anti-FLAG double immunoprecipitation (Re-IP)

293T cells (5 × 10^6^) were seeded in 10 cm dishes. The next day, cells were transfected with Lipofectamine 2000 and FLAG-tagged IFIT1 (8 μg), HA-tagged ubiquitin (8 μg), and equimolar amounts of IFIT3 (wild type or Y438E/L442E) or empty vector plasmids. Six hours post transfection, Lipofectamine was removed, and 48 hours post transfection cells were treated with MG132 for 6 hours, then lysed using lysis buffer (50 mM Tris-HCl pH7.4, 150 mM NaCl, 1 mM EDTA, 1% Triton X-100) supplemented with 1:100 protease inhibitor cocktail (Merck). Samples concentration was measured using a BCA assay (TFS), the total concentration was normalised for all samples and adjusted to 1 mL before immunoprecipitation. Per sample 40 µL of anti-FLAG agarose gel (Merck) was used. Agarose gel was separated from supernatant by centrifugation at 5,000 ×g for 30 seconds at 4 °C followed by 2 minutes incubation on ice to allow the resin to settle (same for all following centrifugations). The supernatant was removed, and the resin was prewashed using 500 µL Tris Buffered Saline (TBS - 50 mM Tris:HCl pH7.5, 150 mM NaCl) twice, then 0.1 M glycine:HCl pH3.5 once, and finally in TBS three times. Samples were then added to the gel and incubated with rotation for 10 minutes at room temperature, then overnight at 4 °C. Supernatant was removed by centrifugation and agarose gel was washed three times using 500 μL TBS with gentle pipetting between washes. Precipitated proteins were eluted by heating samples in 70 µL of ReIP sample buffer (16% glycerol, 1.26% SDS, 4% β-mercaptoethanol, 0.6% Triton X-100, 40 mM Tris pH 7.4, 90 mM NaCl, 0.6 mM EDTA at 80 °C for 10 minutes. The eluates were then diluted 1:10 with fresh protease inhibitor-supplemented lysis buffer and subjected to the second anti-FLAG immunoprecipitation as described above. Samples were then analysed by immunoblotting with indicated antibodies.

### Western blotting

Cells were lysed with RIPA buffer (50 mM Tris-HCl pH8, 150 mM NaCl, 1 mM EDTA, 1% Triton X-100, 0.1% SDS) supplemented with 1:100 protease inhibitor cocktail (Merck). Cell lysates were cleared, and total protein concentration was measured using a BCA assay (TFS) according to the manufacturer protocol. After SDS-PAGE, proteins were transferred to 0.45 μm nitrocellulose membrane (Merck) using Trans-Blot SD semi-dry transfer cell (Bio-Rad).

Membranes were blocked in PBS with 0.1% Tween 20 (PBST) and 5% skimmed milk (Marvel), incubated with primary antibodies diluted in PBST with 2% bovine serum albumin overnight at 4 °C, and incubated for one hour at room temperature with secondary antibodies diluted in PBST before being visualised using the Odyssey CLx Imaging System (LI-COR). Protein bands were analysed using Image Studio Lite 5.2 software.

#### Antibodies

Primaries Anti-FLAG M2 (#F3165, Sigma-Aldrich, 1:1000), anti-IFIT1 (#14769, Cell Signalling Technology, 1:1000), anti-IFIT2 (#12604-1-AP, Proteintech, 1:1000), anti-IFIT3 (#15201-1-AP, Proteintech, 1:1000), anti-PKR (#ab184257, Abcam, 1:2000), anti-HA (#51064-2-AP Proteintech, 1:1000), anti-ISG15 (#15981-1-AP, Proteintech, 1:1000), anti-IFITM1 (#60074-1-Ig, Proteintech, 1:2000) and anti-GAPDH (#60004-1-Ig, Proteintech, 1:8500). Secondaries anti mouse and anti-rabbit (#926-32210, #926-32211, #926-68020, #926-68021, LI-COR, 1:10000).

### Virus infection

293T IFIT1/IFIT3 double KO cells were transfected with 0.4 pmol of each plasmid (using pTwist EF1α constructs). After overnight incubation, cells were infected with SFV6-mCherry virus, MOI 2.5, in media without FBS for 90 minutes, then infection media was removed, and cells were incubated with complete media for 8 hours.

### Flow cytometry

Cell harvesting with trypsin was followed by fixation in 4% paraformaldehyde (TFS). Cell staining was performed in permeabilization buffer (1% FBS, 0.25% saponin in 1X PBS). For the effect in endogenous ISG15 or IFITM1 expression experiments, cells were stained with anti-ISG15 antibody (Proteintech, #15981-1-AP, diluted 1:100) or anti-IFITM1 (#99969, Cell Signaling, diluted 1:100), and anti-IFIT1 antibody (Abcam, #ab118062, diluted 1:100) followed by anti-Rabbit antibody conjugated to Alexa Fluor™ 647 (TFS, #A21244, diluted 1:500) plus anti-Mouse antibody conjugated to Alexa Fluor™ 488 (TFS, #A11001, diluted 1:500). SFV infected cells were stained with anti-IFIT1 antibody (Cell Signaling, #14769, diluted 1:100), followed by anti-Rabbit antibody conjugated to Alexa Fluor™ 647 (TFS, #A-21244, diluted 1:500). Flow cytometry performed in LSRFortessa™ (BD Biosciences) and data analysis in FlowJo (BD Biosciences).

