## Supplementary Info for "IFIT3 controls IFIT1 accumulation and specificity preventing self mRNA targeting during the innate immune response"

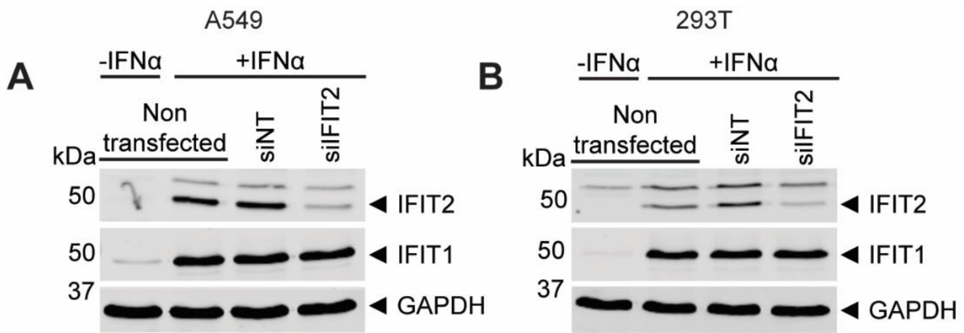


**Supplementary Figure S1** IFIT2 depletion does not impact on IFIT1 levels. A549 (A) and 293T (B) cells were transfected with pooled siRNA for non-targeting (siNT) or IFIT2 (siIFIT2). After 24 hours cells were treated with 100 U/mL of IFNα for a further 24 hours. Cells lysates were collected and analysed for immunoblotting 48 hours post transfection, probing with antibodies for the indicated proteins. GAPDH was used as loading control.

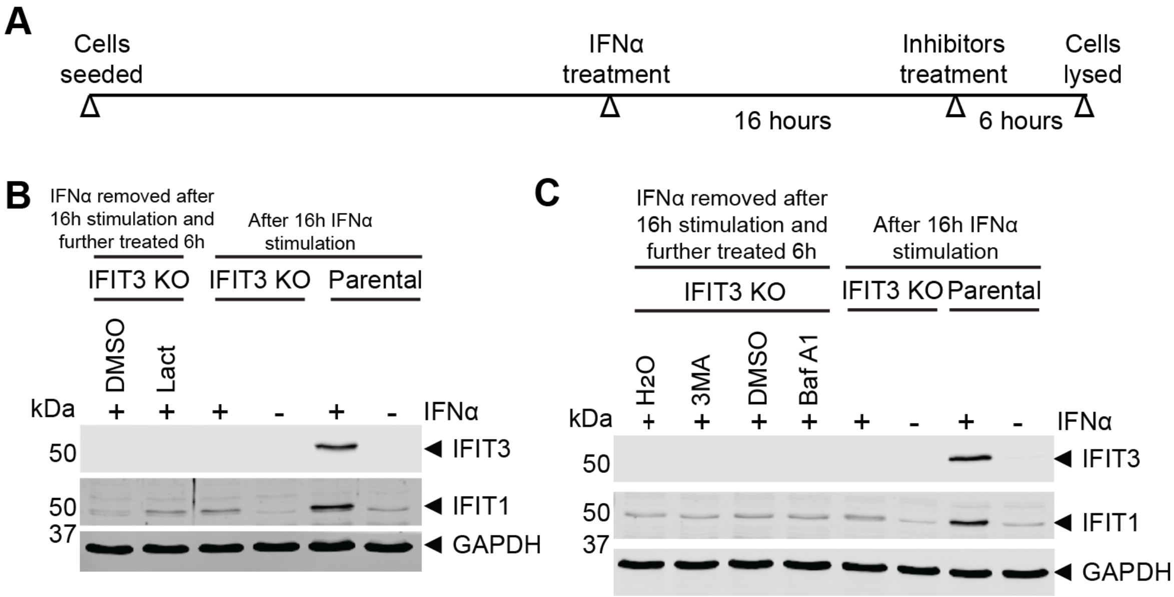


**Supplementary Figure S2** Proteasome, but not autophagy-lysosome, inhibitors rescue IFIT1 levels. (A) Schematic representation of experiment performed in B-C. IFIT3 KO A549 cells were stimulated with 100 U/mL IFNα for 16 hours, then washed twice, and incubated with fresh medium supplemented with (B) lactacystin (Lact), (C) 3-methyladenine (3-MA) or bafilomycin A1 (Baf A1) or vehicle controls for further 6 hours, after which cell lysates were harvested for immunoblotting. Wild type parental cell lysates were included as controls. GAPDH was used as loading control and membranes probed with antibodies for the indicated proteins.

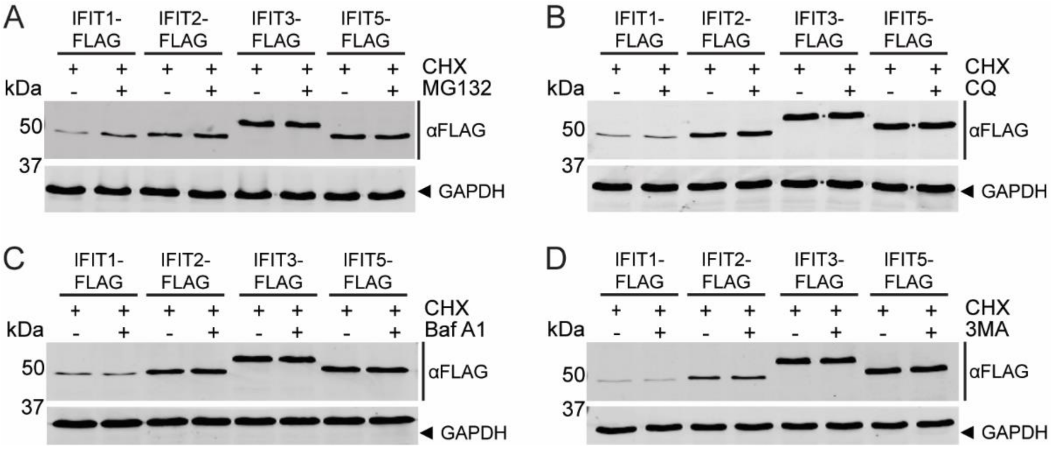


**Supplementary Figure S3** Overexpressed IFIT1-Flag is degraded by the proteasome. Wild type 293T cells were transfected with 1.5 µg of different FLAG-tagged IFITs encoding plasmids for 24 hours and then incubated with fresh medium supplemented with cycloheximide (CHX) plus (A) MG132, (B) chloroquine (CQ), (C) bafilomycin A1 (Baf A1), (D) 3-methyladenine (3-MA) or their vehicle controls for 6 hours. Cell lysates were then extracted for immunoblot analysis and membranes probed with anti-FLAG and anti-GAPDH antibodies. GAPDH was probed as loading control.

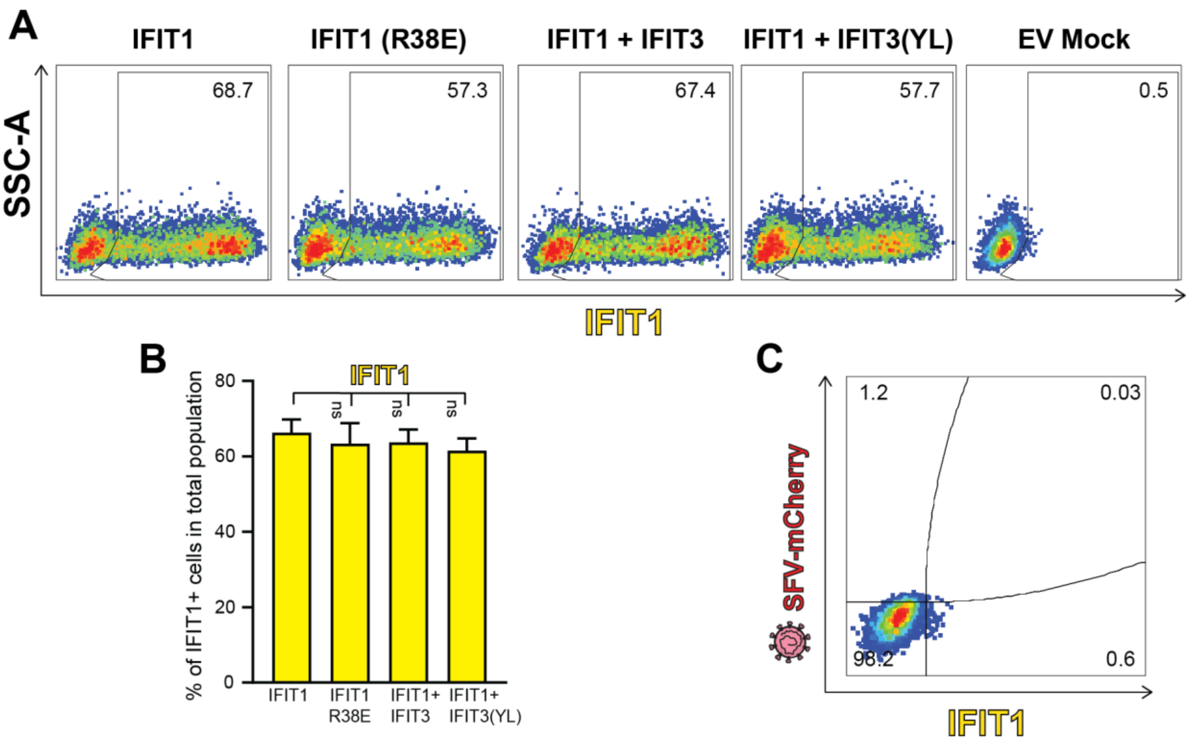


**Supplementary Figure S4** IFIT1 expression levels in SFV infection flow cytometry experiment. A) Gating showing IFIT1 positive cells in total population from representative experiment shown in Figure 4E. B) Quantification of the percentage of IFIT1 positive cells in three experiments (as shown in A). C) Empty vector nontreated control from experiment shown in Figure 4E. SSC-A: Side Scatter Area. In B, data are mean +/- standard deviation (SD) of three independent experiments. Statistical analysis using one-way ANOVA. Asterisks indicate where the percentage of double positive cells was significantly different ($p < 0.05$) between the IFIT1 WT and the other conditions as indicated. ns=not significant.

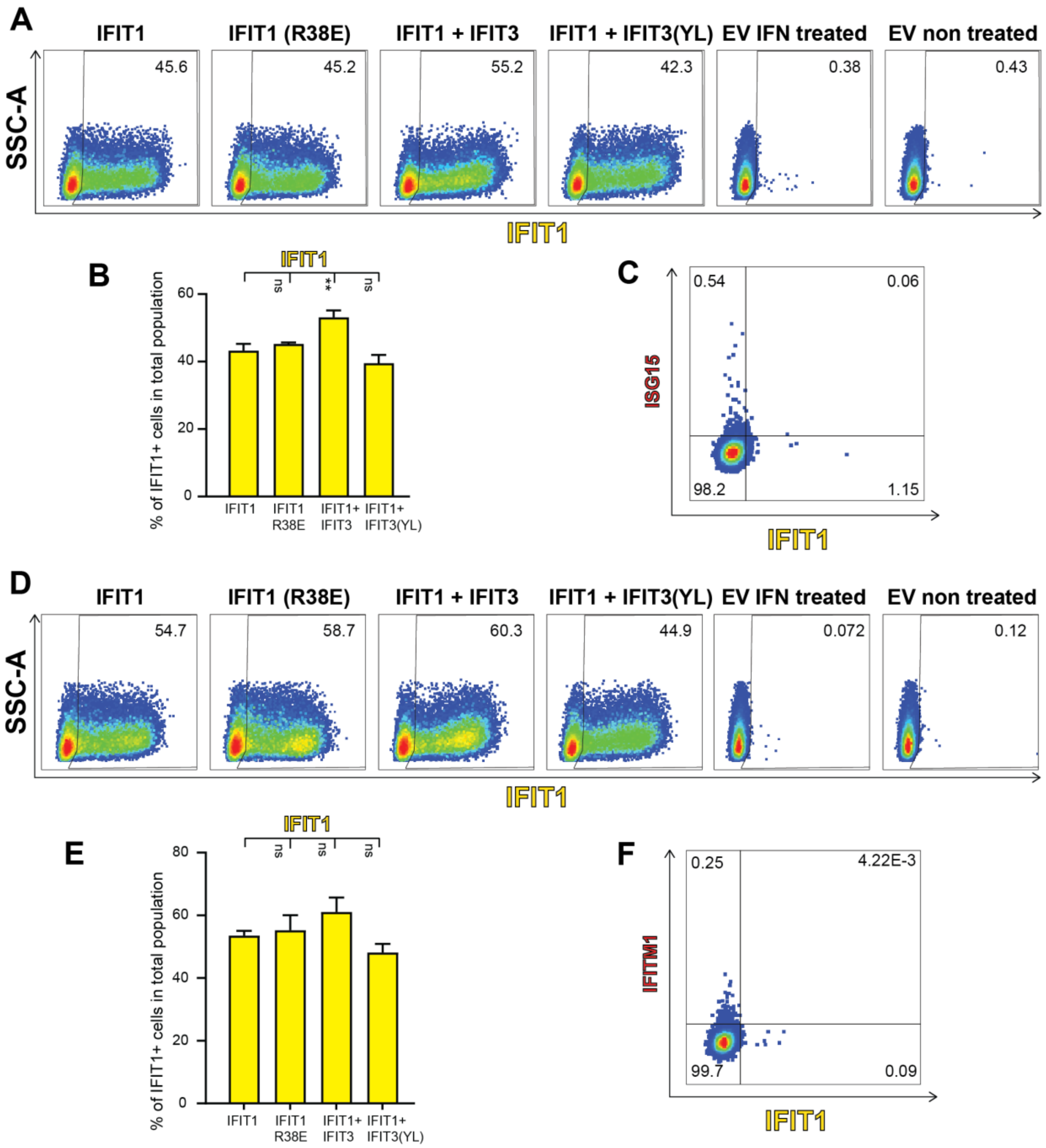


**Supplementary Figure S5** IFIT1 expression levels in ISG15 and IFITM1 flow cytometry experiments. A) Gating showing IFIT1 positive cells in total population from representative experiment shown in Figure 5C. B) Quantification of the percentage of IFIT1 positive cells in three experiments (as shown in A). C) Empty vector nontreated control from experiment shown in Figure 5C. D) Gating showing IFIT1 positive cells in total population from representative experiment shown in Figure 5E. E) Quantification of the percentage of IFIT1 positive cells from three experiments (as shown in D). F) Empty vector nontreated control from experiment shown in Figure 5E. SSC-A: Side Scatter Area. In B and E, data are mean +/- standard deviation (SD) of three independent experiments. Statistical analysis using one-way ANOVA. Asterisks indicate where the percentage of double positive cells was significantly different ($p < 0.05$) between the IFIT1 WT and the other conditions as indicated. ns=not significant.
